# Long reads from Nanopore sequencing as a tool for animal microbiome studies

**DOI:** 10.1101/2019.12.21.886028

**Authors:** Beatriz Delgado, Magdalena Serrano, Carmen González, Alex Bach, Oscar González-Recio

**Affiliations:** Departamento de Mejora Genética Animal, Instituto Nacional de Investigación y Tecnología Agraria y Alimentaria, 28040 Madrid, Spain; Departamento de Producción Agraria, Escuela Técnica Superior de Ingeniería Agronómica, Alimentaria y de Biosistemas, Universidad Politécnica de Madrid, Ciudad Universitaria s/n, 28040 Madrid, Spain; Institució Catalana de Recerca i Estudis Avançats (ICREA), 08010 Barcelona, Spain; Department of Ruminant Production, Institut de Recerca i Tecnologia Agroalimentàries (IRTA), 08140 Caldes de Montbui, Spain

**Author notes:** Correspondence: Oscar González-Recio, Departamento de Mejora Genética Animal, Instituto Nacional de Investigación y Tecnología Agraria y Alimentaria, 28040 Madrid, Spain.

## Abstract

In the era of bioinformatics and metagenomics, the study of the ruminal microbiome has gained considerable relevance in the field of animal breeding, since the composition of the rumen microbiota significantly impacts production and the environment. Illumina sequencing is considered the gold standard for the analysis of microbiomes, but it is limited by obtaining only short DNA sequences to analyze. As an alternative, Oxford Nanopore Technologies (ONT) has developed a new sequencing technique based on nanopores that can be carried out in the MinION, a portable device with a low initial cost which long DNA readings can be obtained with. The aim of this study was to compare the performance of both types of sequencing applied to samples of ruminal content using a similar pipeline. The ONT sequencing provided similar results to the Illumina sequencing, although it was able to classify a greater number of readings at the species level, possibly due to the increase in the read size. The results also suggest that, due to the size of the reads, it would be possible to obtain the same amount of information in a smaller number of hours. However, detection of archaeal and eukaryotic species is still difficult to accomplish due to their low abundance in the rumen compared to bacteria, suggesting different pipelines and strategies are needed to obtain a whole representation of the less abundant species in the rumen microbiota.

## Background

The bovine rumen has been studied for years in an attempt to reveal functions and microorganisms associated with nutritional features such as feed efficiency to use them in animal breeding programs for cattle (Knapp, Laur, Vadas, Weiss, & Tricarico, 2014). Lately, methane emissions from ruminants are among the main concerns in animal husbandry, given their contribution to global warming. Microbial cultures were essential for the first descriptions of the rumen content but they are usually hard to achieve (Creevey, Kelly, Henderson, & Leahy, 2014). Thanks to the Next Generation Sequencing (NGS) techniques it is now possible to detect different microbial taxa in rumen samples avoiding culture (Seshadri et al., 2018). This has allowed analyzing metagenomic samples in an easier way and to detect non-culturable microbes that broaden the knowledge of complex microbial communities as well as allow quantifying the relative abundances of each one of them in the community.

The sequencing by synthesis method utilized by the Illumina platform is currently widely used (Goodwin et al., 2016) and accepted as a gold standard because, among other, its high sequence accuracy. The nanopore sequencing developed by Oxford Nanopore Technologies (ONT) is an attractive technology due to the availability of small and portable devices currently possible at affordable costs (Lu, Giordano, & Ning, 2016). Combined with the diverse kits offered, the ONT is a versatile alternative that can be used for multiple purposes in microbiome studies. There are some publications comparing Illumina and nanopore sequencing (Shin et al., 2016), although most of them focus on 16s rRNA gene sequencing (Cusco et al., 2017; Ma, Stachler, & Bibby, 2017). These previous studies showed a promising potential of nanopore sequencing for species detection with a high correlation between the results obtained by Illumina and ONT at the phylum and genus levels, despite the smaller basecalling accuracy from ONT compared with Illumina. However, latest ONT technology and software developments provide consensus accuracies at genome assembly larger than 99% (Wick, Judd, & Holt, 2019). Nanopore long reads are also being used to assembly whole genomes combining them with short reads for a better accuracy, and the number of microbiome studies using ONT is constantly growing. The clown anemofish, *Amphiprion ocellaris*, has been recently sequenced using hybrid assembly (Tan et al., 2018) as well as a multidrug-resistant COL1 strain (Xia et al., 2017), whose plasmids were also sequenced using short and long reads. This study also showed the ONT potential to make resistome profiles in municipal sewage, obtaining similar results as Illumina, although it was still necessary to improve the throughput of the sequencing runs. ONT has also been used to detected arbovirus in mosquitos (Batovska, Lynch, Rodoni, Sawbridge, & Cogan, 2017) obtaining comparable results to Illumina sequencing, despite the poorer quality of the reads was poorer. Other studies focus on the applicability of ONT for real-time viral pathogen detection (Greninger et al., 2015) in human blood as it shows similar results to Illumina but provides faster results, which is essential in the field of medicine. Moreover, analysis of the 16S rRNA gene is still preferred over shotgun metagenomics using the MinION and most of these studies have been conducted on mouse (Shin et al., 2016) and human gut microbiota (Leggett et al., 2017), proving consistent results for individual taxonomic profiles. Other animal microbiomes have also been sequenced with the MinION, the dog skin microbiota was characterized using a mock community and dog skin samples to test new 16s rRNA primers, finding accurate taxonomic results at the genus level (Cusco et al., 2017).

There is a lack of publications comparing the performance of nanopore sequencing using shotgun sequencing rather than 16s rRNA gene sequencing applied to non-human microbiomes such as the rumen microbiota. Furthermore, to our knowledge, there are still no publications comparing ONT and Illumina to sequence rumen microbiota. Assessing the ONT performance in animal microbiota is relevant as there are several applications for animal husbandry. For instances, nutrition and breeding strategies are being developed to modulate the microbiome in several species (ref). It may be possible to use this technology to detect favorable or pathogen microbial profiles for economically and environmentally relevant traits.

Hence, the objective of this study was to evaluate the performance of ONT using the MinION (Oxford Nanopore Technology, Oxford, UK) device to sequence the rumen microbiota and compare the results with an Illumina MiSeq platform as benchmark in terms of reads yield, taxonomic assignment, and alpha and beta-diversity.

## 2. Methods

### 2.1 Sample collection

Rumen content (50 μL) from 80 animals was extracted using a stomach tube connected to a mechanical pumping unit and was immediately frozen and stored at -80 °C to avoid microbial growth and to preserve the sample from degradation. Samples were later thawed and ground until solid and liquid phases were homogenized using a blender. DNA from 250 μL of each sample was extracted using the “DNeasy Power Soil Kit” (QIAGEN, Valencia, CA, USA). DNA samples of 12 of those animals were selected for extreme phenotype of feed efficiency (FE), calculated as milk production (kg/d) divided by feed consumption (kg/d), 6 for each type of phenotype of feed efficiency. Each sample was analyzed for quality measurement using a Nanodrop ND-1000 UV/Vis spectrophotometer (Nanodrop Technologies Inc., DE, USA), recording their 260/280 and 260/230 ratios.

### 2.2 DNA sequencing

For Illumina sequencing, samples were diluted to a concentration of 5 ng/μL in a total volume of 15 μL/sample in a 96-well plate and were later sequenced using Illumina MiSeq in an external sequencing service (FISABIO, Valencia, Spain). The quality control was performed by the same sequencing service using the prinseq-lite program (Schmieder & Edwards, 2011) and the forward- and reverse-reads from the sequencing were joined using the FLASH software (Magoč & Salzberg, 2011).

The MinION device was used for ONT sequencing. The Ligation Sequencing Kit (SQK-LSK109 for 1D metabarcoding) was used for the library preparation as described by the manufacturer. Reads were basecalled and demultiplexed using Guppy (community.nanoporetech.com). The resulting FASTQ files were used as input of Porechop (Wick R., 2017) for adaptor removal and barcode de-multiplexing. A total of 7,111,651 paired reads were obtained from Illumina sequencing after the QC (Table 1). After the QC for ONT reads, a total of 2,927,404 good quality reads with an average length of 1,885 bp (Table 1) were retained and the distribution of the reads among the barcodes was homogeneous.

**Table 1.** Summary of Illumina and Nanopore sequencing performance and analysis results.

| Sample | Illumina<br>reads | Nanopore<br>reads |
| --- | --- | --- |
| 1147 | 763,635 | 242,794 |
| 1149 | 614,876 | 268,196 |
| 678 | 580,763 | 221,340 |
| 17246 | 519,271 | 229,538 |
| 2631 | 836,541 | 245,231 |
| 152 | 425,134 | 272,455 |
| 8172 | 501,871 | 204,676 |
| 8627 | 784,286 | 273,294 |
| 9524 | 575,278 | 307,658 |
| 8815 | 508,088 | 269,539 |
| 3404 | 534,667 | 147,489 |
| 2954 | 467,241 | 245,194 |
| TOTAL reads | 7,111,651 | 2,927,404 |
| Total yield | 2.13 Gb | 5.48 Gb |
| *Species* | 125 | 146 |
| *Genera* | 78 | 86 |
| *Phyla* | 17 | 21 |
| *Read length*<br>*(pb)* | 2 x 300 | ≈1,885 |

Illumina sequence data are available from NCBI database, with bioproject number PRJNA423103. ONT sequences are available from the authors upon reasonable request.

### 2.3 Data analysis

The FASTQ files were aligned against the NCBI-nr protein database (Nov. 2017) using DIAMOND v0.9.22 (blastx option) setting the -F option to 15, to consider frame-shift errors in the sequences and the – rangeCulling and –top options set to 10 to scan the whole sequence for alignments with a 10% of the best local bit score (megan.informatik.uni-tuebingen.de, accessed on October 2018). The taxonomic binning of short reads from Illumina was performed using the daa2rma program from MEGAN Community Edition (CE) v6.11. with the option -a2t, to map the reads to the NCBI-taxonomy mapping file containing protein accessions (May 2017). Long reads from ONT were analyzed with specific parameters for long reads -lg (long reads) and -alg set to “longReads” (Huson et al., 2018). Relative taxonomic abundances were obtained for each samples and platform representing the number of reads assigned to each taxon. Relative abundances, alpha (Shannon and Simpson indexes) and beta diversity -using Bray-Curtis dissimilarity-were analyzed using the phyloseq (McMurdie & Holmes, 2013), vegan (Oksanen et al., 2019), and microbiome R (Leo Lahti et al., 2012-2019) packages.

### 2.4 Assembly

Short reads from the Illumina dataset were assembled using Megahit (Li, Liu, Luo, Sadakane, & Lam, 2015) with the default parameters using the joined forward and reverse reads. The nanopore reads were assembled using Canu (Koren et al., 2017) setting the options minReadLength = 100, minOverlapLength = 100 as the resulting reads were too short for the default parameters (less than 10,000 bp), and genomeSize = 2.5m, as described in the documentation for metagenome assemblies. The quality of the assemblies was assessed using Quast v4.0 (Mikheenko, Valin, Prjibelski, Saveliev, & Gurevich, 2016).

## 3. Results

### 3.1 Rarefaction curves

Nanopore sequencing had a low throughput in terms of number of reads when compared with Illumina due to the sequencing technology itself. On the other hand, ONT sequencing obtained longer reads as opposed to Illumina. However, rarefaction curves showed that intra-sample diversity was well represented for both technologies (Supplementary Figure 1), suggesting that a low number of reads from nanopore sequencing is enough to represent the microbial complexity in rumen samples. The 48-h sequencing runs in the MinION could be shortened, thus allowing an even faster analysis.

### 3.2 Alpha diversity

Alpha-diversity indexes -Shannon and Simpson-showed a high similarity for each sample in both sequencing datasets, although samples ranked slightly different between platforms (Figure 1). However, the average indexes by platform are highly similar for both the Illumina and ONT datasets -3.20 and 3.30 for the Shannon index respectively, and 0.912 and 0.918 for the Simpson index. The platform effect was not significant (P>0.05) when it was included in a generalized linear model with Shannon or Simpson diversity indices as dependent variable, and parity and platform as explanatory effects.

**Figure 1.**
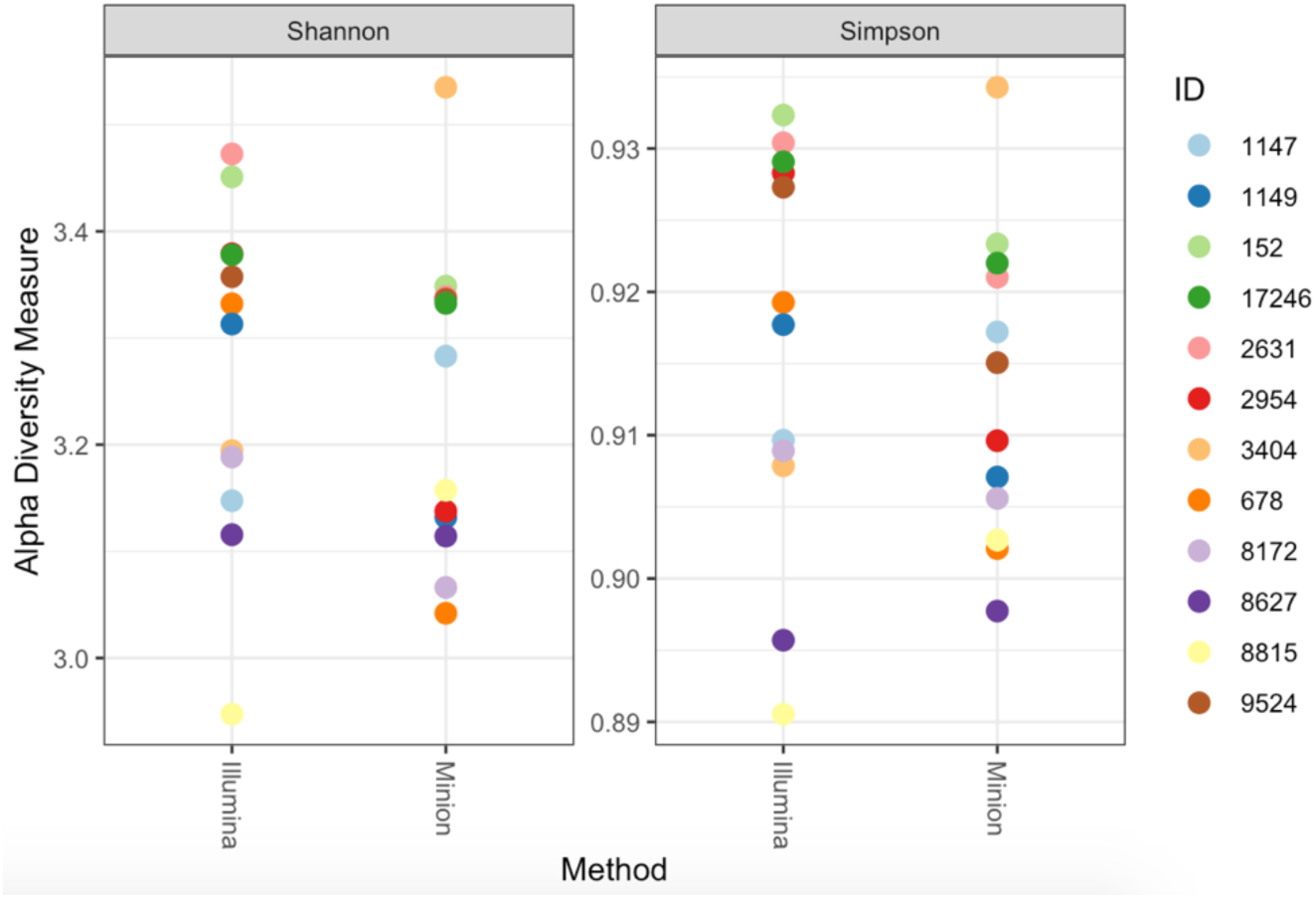
Comparison of alpha diversity indexes. Shannon (left) and Simpson (right) indexes were calculated for each sample and sequencing technology to compare the overall taxonomic diversity.

**Figure 1.**
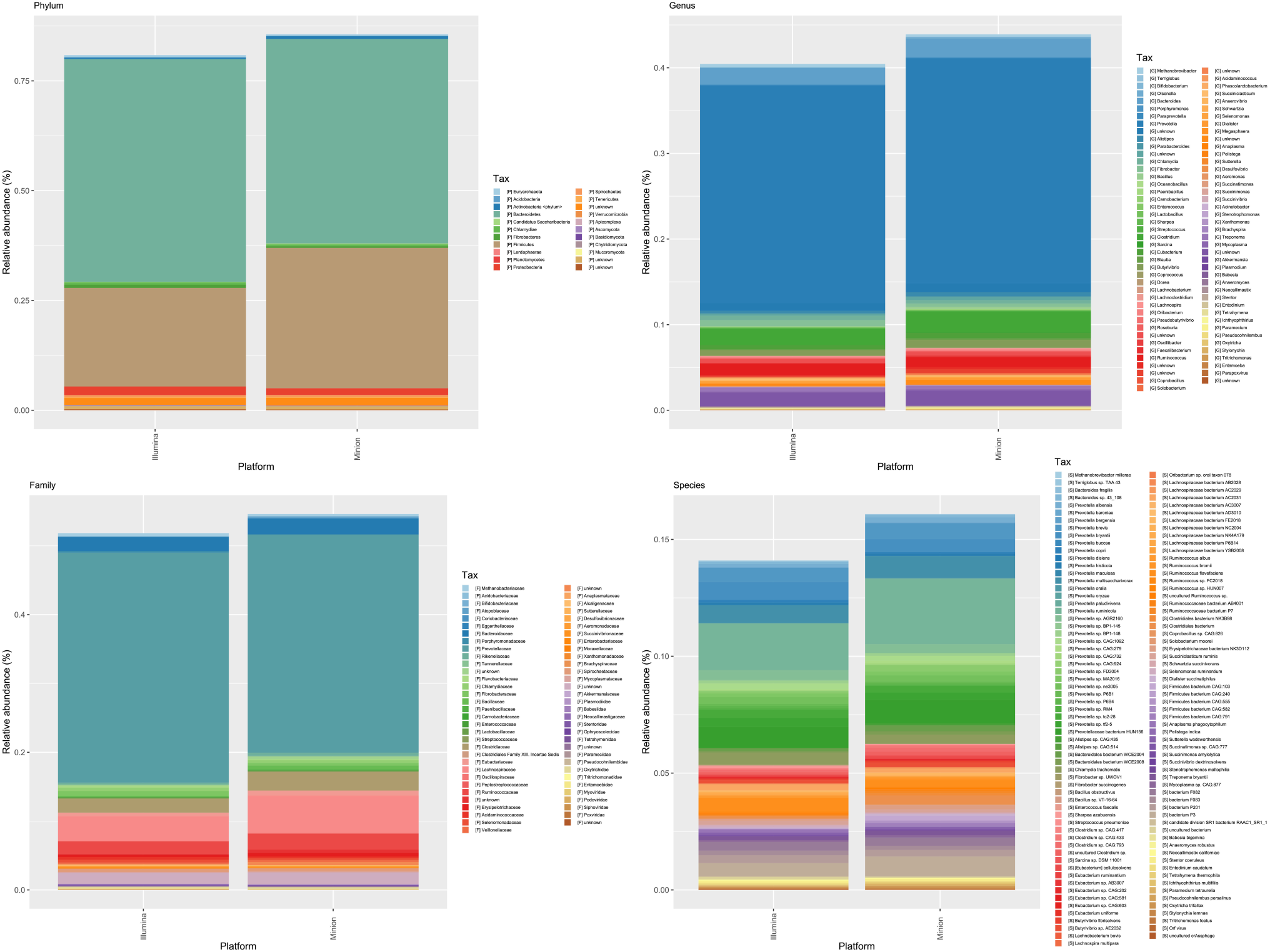
Comparison of relative abundances of common taxa at four different taxonomic levels using Illumina and MinION sequencers. Relative abundances are similar for common taxa at all levels, although the phylum Firmicutes has a higher relative abundance in the MinION dataset which counts for most of the observed differences in the four graphics.

### 3.3 Beta diversity

Nanopore and Illumina sequencing also delivered similar results for microbial composition and abundance at all taxonomic levels, although the nanopore dataset detected a slightly greater diversity (Table 1). Nanopore sequencing seems to detect a greater ecological diversity in the samples at lower taxonomic levels (e.g. species and genera). Longer reads allow DIAMOND and MEGAN assigning more sequences to the reference database as they rely on sequence similarity, and it is more likely to match a read to a species when using longer reads. Both platforms display similar results at upper taxonomic levels -phylum, family and genus-as shown by the similar distribution found in the ordination plots, despite the differences in the number of reads. Inter-sample relative abundances are similar for both sequencing techniques although there were some differences between MinION and Illumina datasets. These differences are greater at the species level and seem to affect sample distribution between both sequencing platforms, meaning the relative abundances of the same sample differ for the different platforms.

### 3.4 Relative abundances

Bacteria, Archaea and Eukarya kingdoms were detected in the expected ratios, with Bacteria being the most abundant taxonomic kingdom. Illumina reads were grouped in 125 species, 78 genera and 17 phyla (Table 1), whereas a larger number of groups were detected from ONT reads: 146 species, 86 genera and 21 phyla. Relative abundances for common taxa were similar, although the MinION detected a greater abundance of the phylum Firmicutes (Figure 3) while maintaining the abundances of other taxa such as Bacteroidetes, which also affects the Bacteroidetes/Firmicutes ratio of each sample (Table 2, Supplementary Figure 2), creating high discrepancies in samples such as number 2631 and 3404. Moreover, a higher diversity was detected with longer reads, since 95 Firmicutes species and 50 Bacteroidetes species could be classified with ONT, whereas when using Illumina reads only yield 72 Firmicutes and 39 Bacteroidetes species respectively could be classified. ONT classified 16% of reads to species level, 2 percentual points larger than Illumina. For comparison, the Bacteroidetes/Firmicutes ratio was analyzed from 16S Illumina sequences (V3-V4 regions). Clear differences were observed in this ratio between Illumina shotgun and Illumina 16S amplicon sequencing (Supplementary Figure 2). The Bacteroidetes/Firmicutes ratio from ONT sequencing overlapped with both Illumina strategies.

**Table 2.**
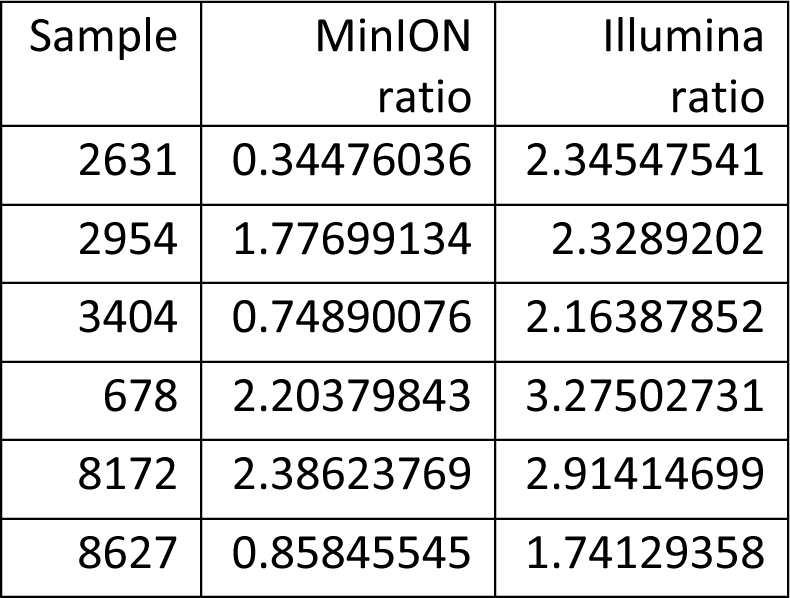
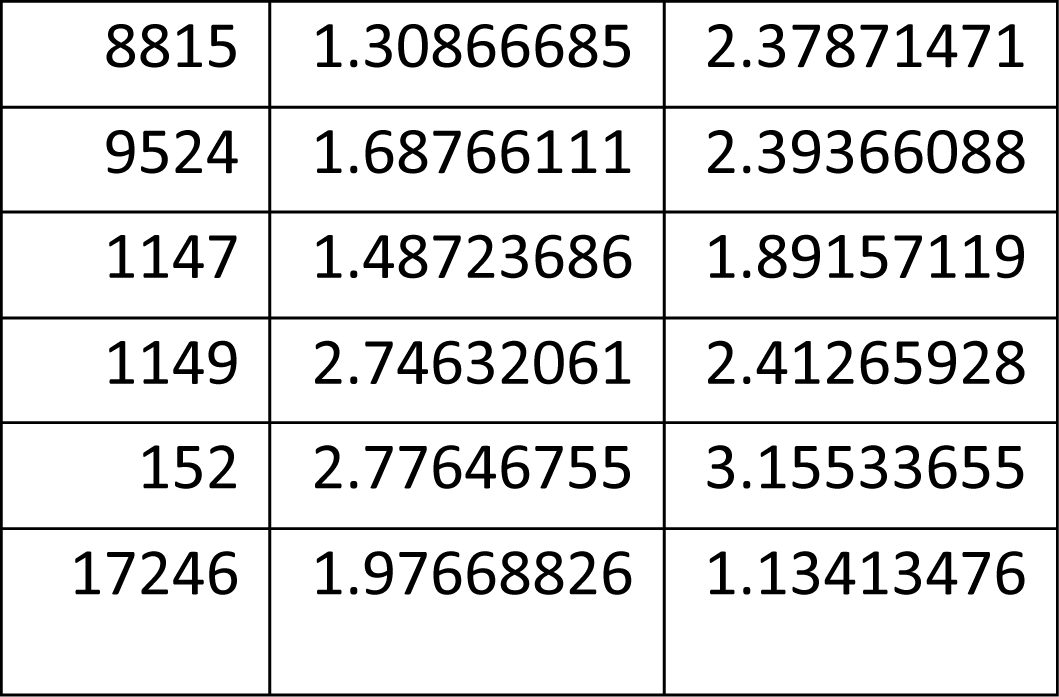
Ratio Bacteroidetes/Firmicutes in both datasets.

| Sample | MinION<br>ratio | Illumina<br>ratio |
| --- | --- | --- |
| 2631 | 0.34476036 | 2.34547541 |
| 2954 | 1.77699134 | 2.3289202 |
| 3404 | 0.74890076 | 2.16387852 |
| 678 | 2.20379843 | 3.27502731 |
| 8172 | 2.38623769 | 2.91414699 |
| 8627 | 0.85845545 | 1.74129358 |
| 8815 | 1.30866685 | 2.37871471 |
| 9524 | 1.68766111 | 2.39366088 |
| 1147 | 1.48723686 | 1.89157119 |
| 1149 | 2.74632061 | 2.41265928 |
| 152 | 2.77646755 | 3.15533655 |
| 17246 | 1.97668826 | 1.13413476 |

**Figure 3.**
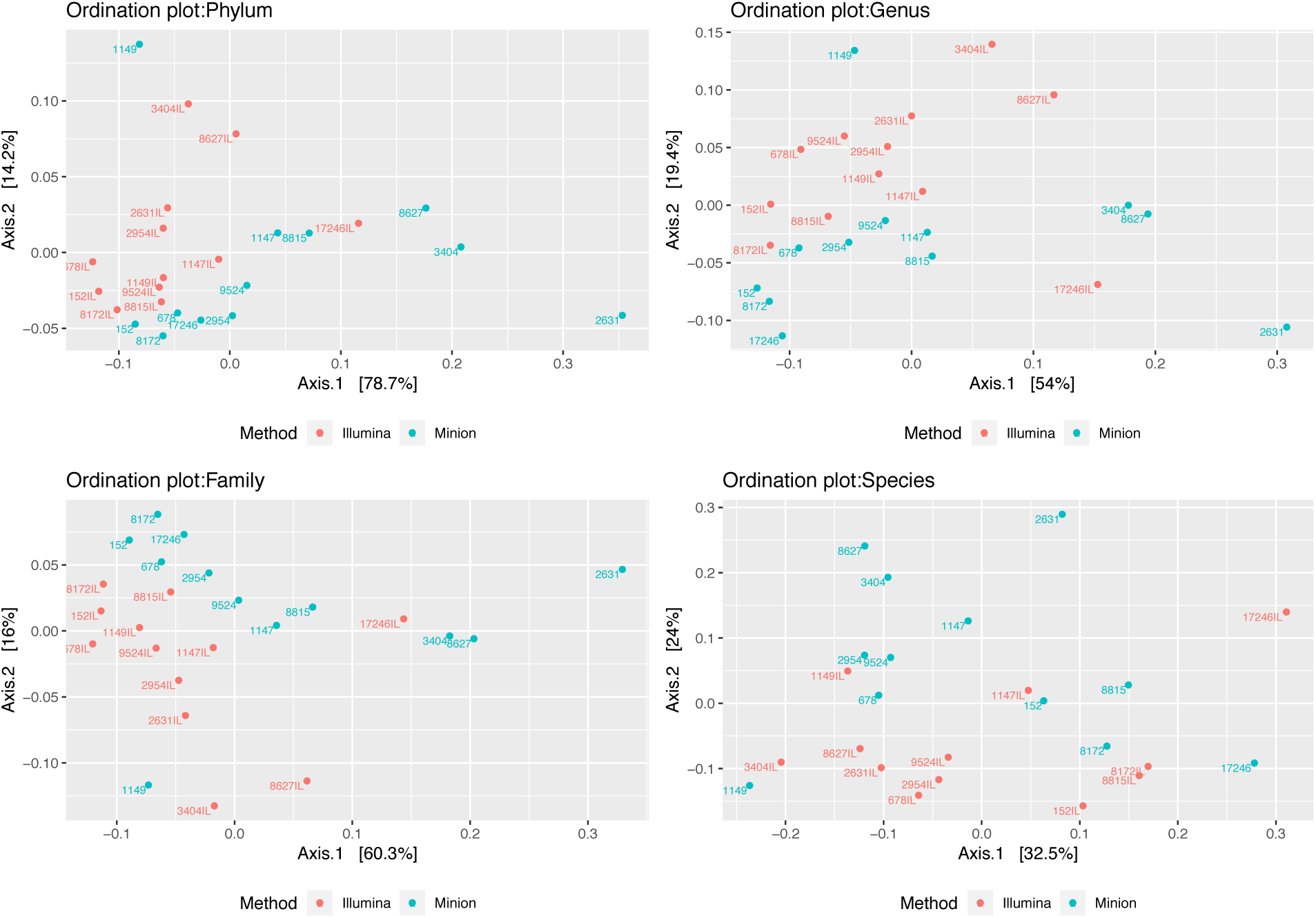
Ordination plots corresponding to four different taxonomic levels. Ordination analysis was performed using data corresponding to relative abundances for both sequencing techniques at different taxonomic levels: phylum, family, genus and species. For the phylum, family and genus levels two different clusters can be observed corresponding to the different sequencing techniques, although sample distribution is similar in both types of sequencing. At the species level no obvious clusters can be observed.

### 3.5 Assembly

Assembly of long and short reads was performed to test the ability of nanopore reads to assembly longer contigs. The assembly of the MinION reads provided a lower number of contigs (281 contigs) compared to the Illumina assembly (7856 contigs), while also having a higher N50 (see Table 3). The total length of the assembly is different -Illumina almost doubles Nanopore’s-which complicates the comparison in terms in the information each assembly provides, but this can be mainly caused by the difference in the number of reads provided by each platform. However, the higher N50 and the low number of contigs from ONT indicate that long reads outperform short reads in metagenomic assemblies, also allowing the assembly of ultra-long contigs -49,178 bp was the longest contigs in the nanopore assembly while Illumina’s longest contig was only 11,173 bp. Sequencing depth with Illumina MiSeq does not seem to be large enough to obtain an assembly with enough quality.

**Table 3.** Assembly statistics.

|  | Illumina assembly | MinION assembly |
| --- | --- | --- |
| Length (bp) | 5,296,436 | 2,550,082 |
| GC (%) | 47.7 | 46.89 |
| N's | 0 | 0 |
| N50 | 641 | 11,549 |
| Largest Contig (bp) | 11,173 | 49,178 |
| Contigs | 7,856 | 281 |

## 4. Discussion

Relative abundance results for common taxa to both platforms were consistent with the previous analyses, finding a high similarity between the two platforms, although the MinION detected a greater abundance of the phylum Firmicutes (Figure 3) while maintaining the abundances of other taxa. Members of the phylum Bacteroidetes have been described as highly efficient polysaccharide degraders due to large genomic regions specialized in this processes (Seshadri et al., 2018) whereas members of Firmicutes are considered “nutritionally fastidious” as they have lost most of their degradative enzymes and feed on fermentation products produced by other microbes. Moreover, the Bacteroidetes/Firmicutes (B/F) ratio has been associated with obesity in some human and mice gut microbiota studies (Koliada et al., 2017), emphasizing that high ratios are related to a higher body mass index (BMI). Obtaining accurate B/F ratios could be key to analyze the relationship between the rumen microbiota composition and relevant phenotypic traits and more efforts are needed to sequence these microbes using nanopore technologies. Other studies in dairy cattle showed that this ratio was strongly associated with milk-fat yield (Jami, White, & Mizrahi, 2014) and suggest that these two phyla could also be related to different feed efficiency parameters such as the dry matter intake (DMI) and residual feed intake (RFI). *Prevotella* spp. -which belongs to Bacteroidetes-has been associated to feed efficiency (Jewell, McCormick, Odt, Weimer, & Suen, 2015; Bach et al., 2019), although most of these results were not consistent due to the low taxonomic resolution of 16S rRNA gene sequencing. Species among this genus are involved in central rumen processes such as pectin and xyloglucan degradation and acetate production (Seshadri et al., 2018), which have a high impact on the nutrition of ruminants. Some previous studies showed that *Prevotella bryantii -*also detected by the two platforms-has some relationship with feed efficiency (Elolimy, Arroyo, Batistel, Iakiviak, & Loor, 2018). The similarities observed for the principal taxonomic groups imply that ONT reads could be used in association analyses to correlate the microbiome composition to phenotypic traits such as feed efficiency. A correlation analysis between the rumen microbiome and feed efficiency has already been performed using Illumina reads, where significant correlations between a high feed efficiency were found when the microbiome composition had a high abundance of Bacteroidetes and a low abundance of archaea (Delgado et al., 2019), which the current study found in almost the same proportion in both datasets. Moreover, the higher number of species classified by long reads indicates that it is likely that nanopore reads are also suitable for other applications such as pathogen detection in cattle since its higher species resolution.

The overall diversity of methanogens assigned by the ONT reads was low, detecting only two species belonging to the genus *Methanobrevibacter*. The species *M. millerae* was the only common species between the two platforms, while the Illumina reads also included other important species such as *M. ruminantium* and *M. olleyae. Methanobrevibacter spp*. In addition, relevant ciliates appear in similar proportions in both datasets, such as *Entodinium caudatum, Tetrahymena thermophila, Paramecium tetraurelia* and *Oxytricha trifallax*, although the presence of these microorganisms was not consistent since some samples had no eukaryotic representatives, which happened both in the ONT and the Illumina dataset. The low detection of important archaeal species such as *M. ruminantium* and *M. olleyae* are one of the most relevant differences between the two platforms. *Methanobrevibacter spp*. is usually the main methanogens in rumen samples and several studies are consistent in the presence of *M. millerae* -which was found in both datasets- and *M. ruminantium* (Chaucheyras-Durand & Ossa, 2014). Moreover, the rumen methanogen *M. millerae* M9 has been shown to have more copies of methanogenesis-related genes (Kelly et al., 2016) and some members of the class Thermoplasmata -also detected by the Illumina reads-have been proposed as a target for methane reduction (Poulsen et al., 2013) as they are methylotrophic and do not require hydrogen to synthesize methane. The ciliates detected by both platforms -*Entodinium caudatum, Tetrahymena thermophila, Paramecium tetraurelia* and *Oxytricha trifallax-*,are also commonly found in the rumen microbiome (Newbold, De la Fuente, Belanche, Ramos-Morales, & McEwan, 2015). Ciliates and methanogens are usually symbionts and are involved in methane production, although only methanogens produce methane (Newbold et al., 2015). Members of Ciliophora have more hydrogenosomes than other protozoa and are believed to have more endosymbiotic methanogens than other protozoa and to produce a greater impact on methanogenesis (Newbold et al., 2015). However, presence of eukaryotic microorganisms was not detected in some samples, which can be due to the low abundance of this species, compared with bacteria in the rumen microbiota or due to sensitivity issues in the pipeline. These results suggest that the taxonomic classification is comparable to that of Illumina at the phylum, family and genus levels, although long reads can be useful for species detection (e.g. pathogen detection). More sensitive procedures, such as amplicon or operon sequencing, may be necessary for a better detection of low abundant microorganisms. Increasing the sequencing depth could also be useful to detect a higher number of non-abundant species (Sims, Sudbery, Ilott, Heger, & Ponting, 2014) such as archaea and eukaryotes in rumen samples. Better extraction methods that avoid DNA fragmentation may improve the number of long-DNA molecules that are recovered from the samples, which also allows a better nanopore sequencing.

Assembly of metagenomic data is the best strategy to recover genomes from the data, while non-assembly strategies are usually better to analyze microbiome profiles. Moreover, assemblies are useful for functional profiling and for connecting relevant metabolic pathways to their corresponding microbes and for discovering and characterizing important species. Although this study suggests that nanopore sequencing outperforms short-read assemblies for more accuracy in the final assemblies, a combination of both types of reads is usually the best way to obtain high quality genomes (Tan et al., 2018), as Illumina’s short reads are usually highly accurate. However, nanopore reads are expected to reach the same accuracies as new bioinformatic tools are being developed so high-quality assemblies could be obtained using only nanopore reads, although the sequencing depth plays an important role in the quality of the assemblies, and it should be increased to obtain better results. The average size of the reads obtained by the MinION must be longer for better assemblies, as they would cover larger parts of the genomes. For that, the protocols for DNA extraction should be improved to avoid shearing the microbial DNA.

Classical approaches for modulating the microbiome include management practices and food additives, genetic selection is the principal strategy to obtain animals with improved phenotypic traits (Tapio, Snelling, Strozzi, & Wallace, 2017). It has been suggested that the microbial composition in ruminants could be used as a predictor of complex traits (Gonzalez-Recio, Zubiria, García-Rodríguez, Hurtado, & Atxaerandio, 2018), as there are signs of host genetic control over the microbiome composition in cows, which could lead to animal breeding programs that select animals with a favorable microbiome for a high feed efficiency phenotype (Sasson et al., 2017). Thus, selecting animals with less methane production could lead to a decrease in the overall impact of livestock in our environment.

## Conclusions

Illumina sequencing is the most used sequencing technology for shotgun metagenomics as it has a high sequencing depth and accuracy, but the short length of the reads makes it difficult to assign specific taxa or genes without a previous assembly. Nanopore sequencing provides long reads that could cover larger genome regions allowing more accurate assemblies. However, long reads involve greater computational efforts. Here we used DIAMOND and MEGAN tools which can work with long reads. Our analyses suggest that ONT sequencing provide an interesting alternative for microbiome profiling in rumen samples following this protocol. The microbiome compositions determined using the two technologies are similar, especially at the phylum and family levels, although a larger number of taxonomic groups is detected when using longer reads. Both platforms provide similar beta-diversity between sequenced samples. The ONT sequencing provides an easier approach to microbiome profiling for almost all kind of laboratories due to its small size, flexibility, and relative low cost. The ONT could even be used in the field to analyze microbiome profiles in real time and provide quick results that may be used for diagnostic or phenotyping purposes.

## Supplementary Figures

**Supplementary figure 1.**
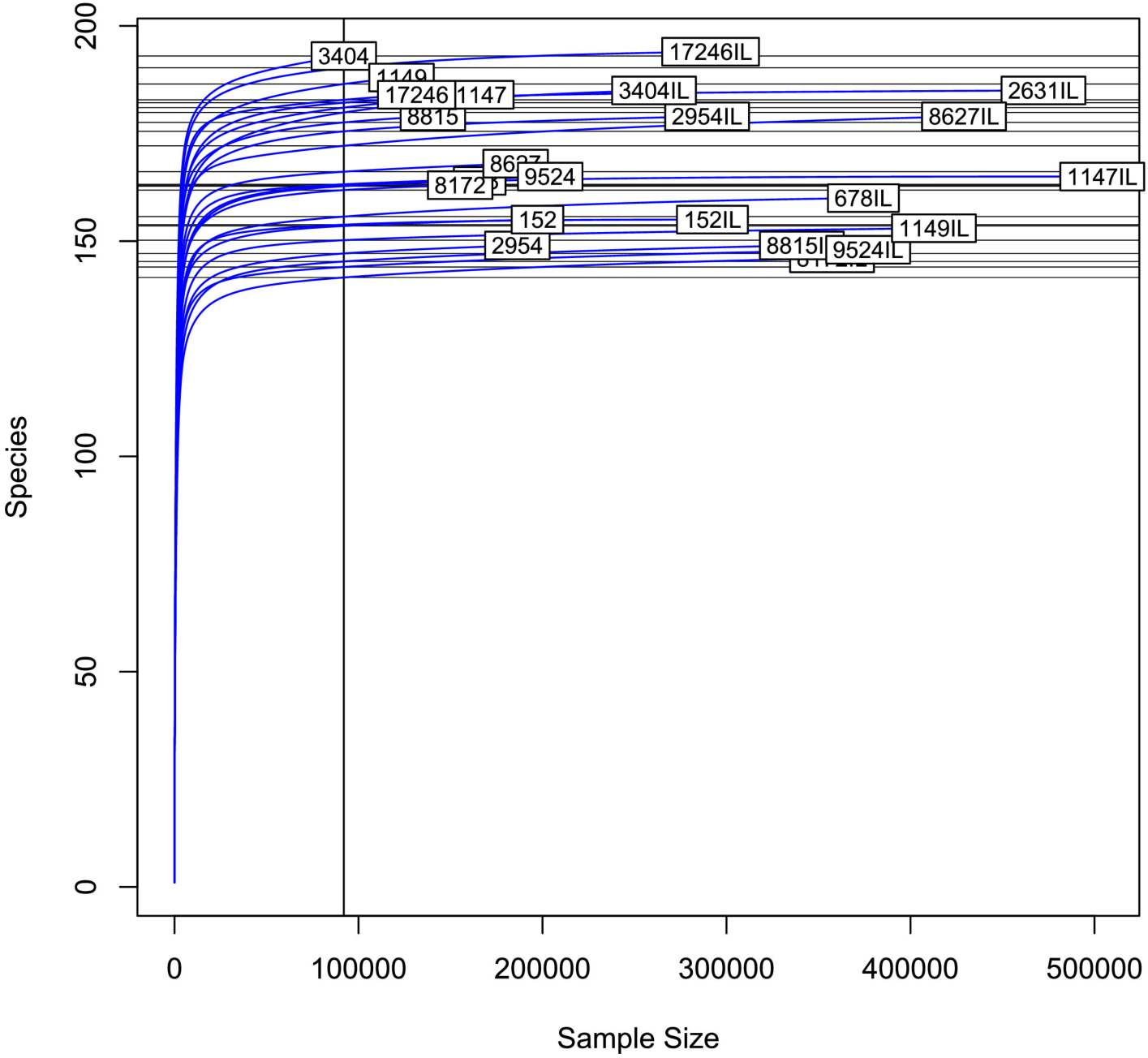
Rarefaction curves of the 12 samples for both sequencing techniques.

**Supplementary figure 2.**
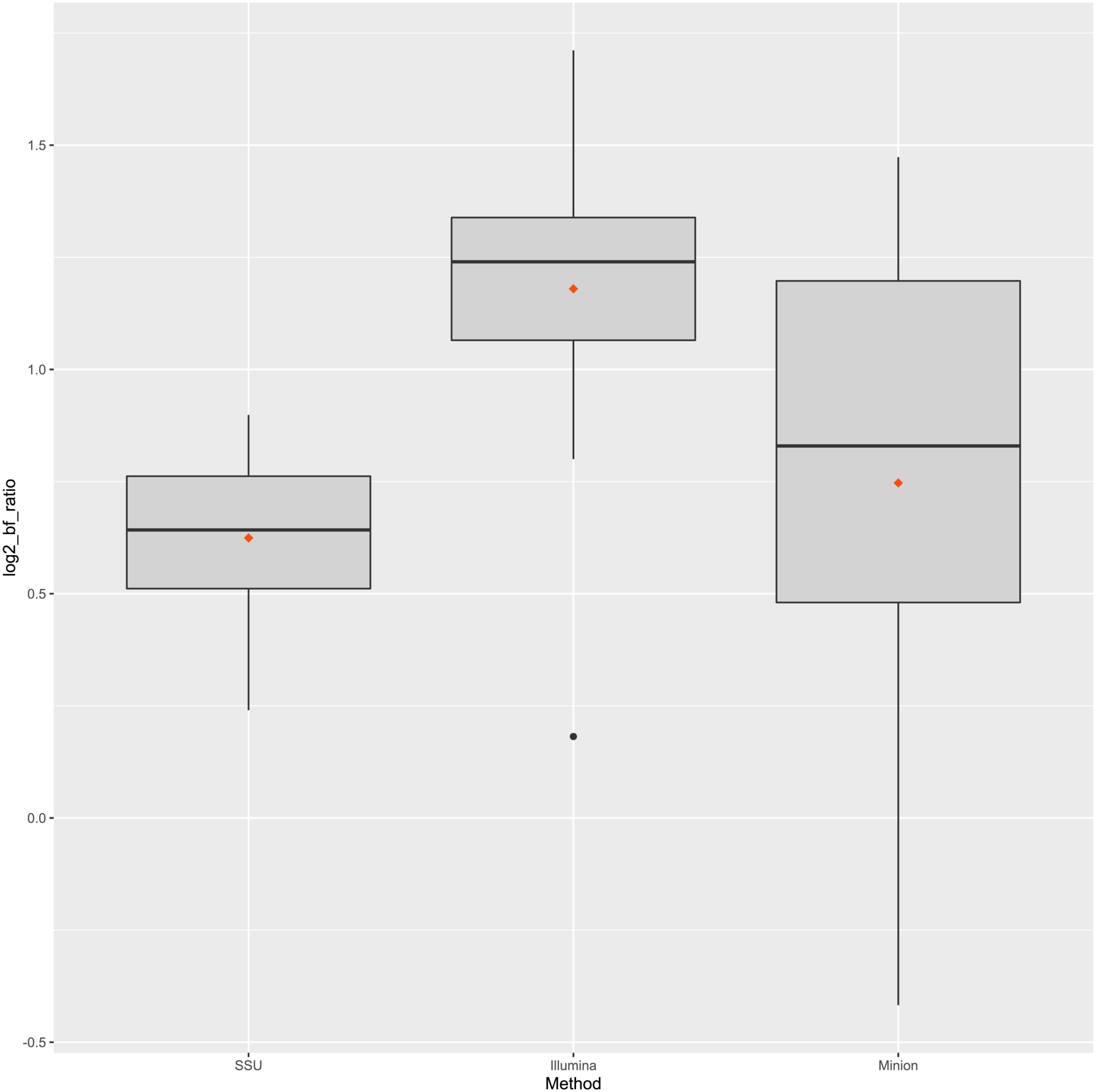
Average ratio Bacteroidetes/Firmicutes per platform.

